# Brief Report: Test-Retest Reliability of Explicit Auditory Processing Measures

**DOI:** 10.1101/2020.06.12.149484

**Authors:** Kazuya Saito, Hui Sun, Adam Tierney

## Abstract

In this brief report, we examined the test-retest reliability of our in-house explicit auditory processing measures in the context of 30 L1 and L2 English users. The participants took the same test battery which consisted of a total of four discrimination tasks (encoding acoustic details of formant, pitch, duration, and rise time) and two reproduction tasks (repeating novel melodic and rhythmic patterns) at Days 1 and 2. According to the results, the participants’ initial and second test scores demonstrated medium-to-large associations (*r* = .562-.907). The results suggest that the tests can tap into various dimensions of individuals’ auditory acuity and integration abilities.

## Introduction

Auditory processing is defined as one’s domain-general capacity to encode, remember, and integrate frequency and time characteristics of sounds, and considered to comprise one dimension of lower-level perception skills (relative to cognitive skills). As basic auditory perception ability is involved in every stage of language learning (e.g., filling in segmental details, detecting prosodic patterns, and identifying word and syntactic boundaries), it has been proposed that individual differences in auditory processing precision serve as a bottleneck for first language (L1) acquisition (Goswami, 2015).

In the cognitive psychology literature, auditory processing has been measured behaviourally through a battery of psychoacoustic tests, wherein learners are *explicitly* asked to encode acoustic details of sounds (i.e., acuity ability), and integrate such auditory information into motor action (i.e., integration ability) (Moore, 2012). Following this methodological standard, our research team at University College London and Birkbeck, University of London has developed (a) four discrimination instruments for acuity, and (b) two different reproduction instruments for integration ability (see Table 1). Whereas the tests are generally delivered to participants at a face-to-face meeting under laboratory conditions, we have begun to implement the test via the online psychology experiment platform, GORILLA (Anwyl-Irvine, Massonnié, Flitton, Kirkham, & Evershed, 2020).

**Table 1.** Constructs of Explicit Auditory Processing and Its Measures.

| Type of processing | Measures |
| --- | --- |
| Auditory acuity | Duration discrimination |
|  | Amplitude rise time discrimination |
|  | Pitch discrimination |
|  | Formant discrimination |
| Audio-motor integration | Rhythm reproduction |
|  | Melody reproduction |

In our research projects involving 300+ second language (L2) English users with a range of L1 backgrounds (Chinese, Japanese, Polish, Spanish) in various contexts (immersion, classroom), auditory acuity scores have been found to account for a medium-to-large amount of variance in L2 comprehension skills (vowel perception, grammaticality judgements) (e.g., Kachlicka, Saito, & Tierney, 2019) and L2 production skills (segmental, suprasegmental, lexical, collocational, and morphosyntactic accuracy) (e.g., Saito, Kachlicka, Sun, & Tierney, 2020; Saito, Macmillan, Kroger, Magne, Takizawa, Kachlicka, & Tierney, forthcoming). Similarly, auditory integration scores have shown significant associations with L2 vowel and prosodic perception proficiency (e.g., Sun, Saito, & Tierney, forthcoming) and L2 phonological and lexicogrammar production proficiency (e.g., Saito, Tran, Suzukida, & Tierney, forthcoming).

In this brief report, we present the results of our first attempt to examine the test-retest reliability of our in-house auditory processing test batteries. The current dataset was collected from 30 English users who took the tests online twice over two consecutive days (T1, T2). Our research question was: Whether and to what degree do their T1 and T2 scores relate to each other? Given that the tests were designed to tap into participants’ auditory acuity and integration abilities, our prediction was that participants’ initial and second test scores would demonstrate significant associations, and the T1/T2 performance would not substantially differ. Note that the test-retest reliability project is ongoing. As more data is collected through our future projects (from June 2020 onward), we plan to expand and update the results.

## Method

### Participants

A total of 30 L1 and L2 English users (13 males, 17 females) participated in the current reliability project, including native speakers of Chinese (*n* = 8), Japanese (n = 8), English (*n* = 5), Polish (*n* = 4), French (*n* = 2), and other Indo-European languages (*n* = 3). Their chronological age ranged from 22 to 49 (*M* = 29.9 years; *SD* = 6.9). 16 out of 30 participants reported music training experience (*M* _*length*_ = 9.0 years, *SD* = 5.8) with varied ages of onset (*M* = 12.8 years, *SD* = 10.3). While eight participants were in English-as-a-Foreign-Language contexts (Japan or China), 27 participants were residing in the UK.

As for L2 English users (*n* = 25), their daily L2 use widely varied (*M* = 53.2%, *SD* = 34.2). Although we did not conduct any L2 English proficiency tests, all of the participants had many years of L2 English learning experience in various settings (classroom, immersion), and they had opportunities to use L2 English for work- and study-related purposes at the time of the data collection. Thus, it is reasonable to assume that the participants in the current dataset were likely to fall into Independent to Proficient Users (but not Basic Users) as per the Common European Framework of Reference for Language.

### Procedure

Due to the ongoing COVID-19 situation, all the data collection was conducted via the online psychology experiment platform, GORILLA (Anwyl-Irvine et al., 2020). Each participant received from a researcher detailed instruction on the main objective of the study, and the procedure for the explicit auditory processing tests. One day after the initial test sessions, participants completed the second test sessions. In each session, the tests were presented in the following order: (1) formant discrimination, (2) pitch discrimination, (3) duration discrimination, (4) rise time discrimination, (5) rhythm reproduction, and (6) melody reproduction. In the following sections, we provide a brief description of the discrimination and reproduction tests.

### Discrimination Tests

The test battery comprised four sub-tests, each of which was designed to assess thresholds for discrimination of differences in a particular acoustic dimension embedded in complex tones. In each sub-test, 100 stimuli were prepared which differed in formant frequencies (F2 = 1500-1700 Hz), pitch (F0 = 330-360 Hz), amplitude rise time (15-300ms), and duration (250-500ms). For a summary of the standard vs. control stimuli, see Table 2. During each trial, three stimuli were presented. Either the first or the third sound was different from the other two. Participants were asked to indicate which sound was different by either pressing the number ‘1’ or ‘3’ on a keyboard. An adaptive three-alternative forced-choice procedure was used. The difficulty of the task decreased after every incorrect response and increased after every third correct response. The program continued until eight reversals were reached, i.e. incorrect answers after a string of successes or correct answers after a string of failures. The threshold was then calculated as the average of the levels of each reversal from the second onward. The scores ranged from 0 to 100 points, indexing how small differences participants could perceive and discriminate. For more methodological details, see Kachilicka et al. 2019; all the synthesized stimuli are available in IRIS. For the subsequent analyses, we used subcomponent scores (formant, pitch, duration, rise time) and overall scores (standardized and averaged scores).

**Table 2.** Summary of Stimuli in Discrimination Tasks.

| Measures | Standard stimuli | Control stimuli | Step size |
| --- | --- | --- | --- |
| Duration discrimination | 250ms | 252.5-500ms | 2.5ms |
| Risetime discrimination | 15ms | 17.8-300ms | 2.8ms |
| Pitch discrimination | 300Hz | 330.3-360Hz | 0.3Hz |
| Formant discrimination | 1500Hz | 1502-1700Hz | 2Hz |

### Reproduction Test

Building on Tierney et al. (2017), participants were asked to listen, remember, and repeat a sequence of complex tones which varied in in fundamental frequencies (for melody reproduction) and in the ratio of drum hits and rest (for rhythm reproduction). For melody reproduction, there were a total of 10 melodies (300 ms per note). At the beginning of the test, participants were shown a set of five buttons, which were arranged in a line stretching from the top to the bottom of the screen. They were encouraged to try clicking on the buttons; pressing each button caused the production of a different tone, with higher frequency tones linked to buttons that were closer to the top of the screen. During the test, for each trial, they were asked to listen to the same sequence repeated three times, and then repeat it by pressing the five buttons. The accuracy ratio (%) was recorded based on fthe irst seven button presses. For rhythm reproduction, a total of 30 rhythmic patterns were generated from Povel and Essens’s (1985) notion of strongly vs. weakly metrical sequences (3.2 seconds per token). After listening to the stimuli, participants reproduced the rhythm by repeatedly pressing the space key. The inter-press times were first quantized by changing them to the nearest interval in the set [200 400 600 800 1000 1200 1400 1600 1800 2000] me. Accuracy ratio (%) was then recorded in terms of the presence of hits or rests at every 200 ms. For more methodological details, see Sun et al., forthcoming; all the melodic and rhythmic sequences are available in IRIS. For the subsequent analyses, we used subcomponent scores (melody, rhythm) and overall scores (standardized and averaged scores).

## Results

### Constructs of Discrimination Test Scores

As summarized in Table 3, the participants’ auditory acuity abilities widely varied in terms of formant, pitch, duration and rise time discrimination. A visual inspection of the data indicated moderate positive skewness especially for the participants’ discrimination performance. According to the results of the Kolmogorov-Smirnov test, whereas the participants’ overall, melody, and rhythm reproduction scores were found to be comparable to normal distribution (*p* > .05), their overall, formant, pitch, duration, and rise time discrimination scores at T1 and T2 significantly differed from normal distribution (*p* < .05). The overall, formant, pitch, rise time, and duration discrimination scores at T1 and T2 (moderate positive skewness) were transformed via a sqrt (x+1) function. For the rest of the Results section, transformed scores were used for the analyses of overall, formant, pitch, duration, and rise time discrimination; and raw scores were used for the analyses of overall, melody, and rhythm reproduction.

**Table 3.** Summary of Raw Auditory Processing Scores.

|  | T1 |  |  | T2 |  |  |
| --- | --- | --- | --- | --- | --- | --- |
|  | *M* | *SD* | *Range* | *M* | *SD* | *Range* |
| Overall discrimination | .00 | .65 | -.77 -1.89 | .00 | .68 | -.95-1.82 |
| Formant discrimination | 46.9Hz | 38.5 | 8.0-193.2 | 38.3Hz | 28.3 | 8.8-132.4 |
| Pitch discrimination | 5.1Hz | 4.0 | 0.9-16.4 | 3.7Hz | 02.8 | 0.8-10.5 |
| Duration discrimination | 48.3ms | 35.7 | 5.6-179.5 | 44.5ms | 25.5 | 6.9-98.5 |
| Rise time discrimination | 58.0ms | 60.1 | 9.8-229.6 | 50.9ms | 54.7 | 6.2-173.6 |
| Overall reproduction | .00 | .89 | -1.56-1.63 | .00 | .90 | -2.08-1.34 |
| Melody reproduction | 73.2% | 23.4 | 30.0-100.0 | 81.6% | 17.9 | 43.0-100.0 |
| Rhythm reproduction | 74.7% | 10.8 | 52.0-98.0 | 80.2% | 10.5 | 59.0-99.0 |

### Constructs of Auditory Discrimination and Reproduction Tests

To check how discrimination and reproduction scores are related to each other on a broad level, overall discrimination and reproduction scores at the outset of the project (T1) were submitted to the Pearson correlation analyses. The results found a weakly significant relationship, *r* = −.377, *p* = .040, indicating that the overlap between the constructs of the participants’ overall discrimination and reproduction abilities could be minor at best.

We ran another set of correlation analyses to further examine the inter-relationships among the subcomponents of discrimination (formant, pitch, duration, rise time) and reproduction tasks (melody, rhythm) at T1. An alpha level was set to *p* < .010 (Bonferroni corrected). According to the results summarized in Table 4, it was unsurprising that the two reproduction (melody, rhythm) demonstrated strong associations (*r* = .618, *p* < .001). When it comes to discrimination, we did not find any significant correlations among the participants’ pitch, duration and rise time scores (*p* > .010); but formant discrimination was significantly linked to pitch discrimination (*p* = .010), rise time discrimination (*p* = .009), and rhythm reproduction (*p* < .001). The results here suggest that the formant component of the discrimination tasks could tap into L2 learners’ composite auditory acuity and integration abilities.

**Table 4.** Inter-Relationships Between Subcomponents of Auditory Acuity and Reproduction Tests.

|  | Pitch discrimination |  | Duration discrimination |  | Rise time discrimination |  | Melody reproduction |  | Rhythm reproduction |  |
| --- | --- | --- | --- | --- | --- | --- | --- | --- | --- | --- |
|  | *r* | *p* | *r* | *P* | *r* | *p* | *r* | *p* | *r* | *p* |
| Formant discrimination | .462* | .010 | -.072 | .704 | .466* | .009 | -.385 | .035 | -.639* | < .001 |
| Pitch discrimination |  |  | .010 | .960 | .322 | .083 | -.457 | .011 | -.353 | .056 |
| Duration discrimination |  |  |  |  | .217 | .249 | .369 | .045 | .001 | .999 |
| Rise time discrimination |  |  |  |  |  |  | -.020 | .915 | -.380 | .038 |
| Melody reproduction |  |  |  |  |  |  |  |  | .618* | < .001 |

*Note.* \* indicates $p < .010$ (Bonferroni corrected)

### Test-Retest Reliability

To check if the behavioural tasks assess a stable construct of one’s auditory processing abilities, the relationship between the participants’ overall and subcomponent test scores at T1 and T2 was examined via a set of Pearson correlation analyses (an alpha level set to *p* < .007, Bonferroni corrected). As shown in Table 5, the participants’ T1 and T2 scores demonstrated significant test-retest correlations, except for duration (*r* = .284, *p* = .128). To check the presence of training effects (T1 to T2 gains), a set of paired sample t-tests were performed as well. Whereas there was no significant difference in the participants’ discrimination scores between T1 and T2, they significantly improved their reproduction scores through taking the test twice (*p* < .001).

**Table 5.** Results of Test-Retest Reliability.

|  | *Correlation analysis* |  | *Paired-sample t tests* |  |
| --- | --- | --- | --- | --- |
|  | *r* | *p* | *t* | *p* |
| Overall discrimination | .701* | < .001 | 0.267 | .791 |
| Formant | .619* | < .001 | 1.440 | .160 |
| Pitch | .562* | .001 | 2.177 | .038 |
| Duration | .284 | .128 | 0.421 | .677 |
| Rise time | .798* | < .001 | 1.406 | .170 |
| Overall reproduction | .863* | < .001 | -5.170* | < .001 |
| Melody | .907* | < .001 | -4.419* | < .001 |
| Rhythm | .775* | < .001 | -4.044* | < .001 |

*Note.* \* indicates $p < .007$ (Bonferroni corrected)

### Factors Affecting Auditory Acuity and Integration

The final objective of our statistical analyses was to examine which learner factors could mediate the degree of individual variation among the participants’ auditory acuity and integration abilities. In this analysis, the participants’ discrimination and reproduction scores at T1 and T2 were averaged and used as dependent variables, respectively. The learner factors were coded as Group, including the participants’ chronological age (*n* = 10 for 20s*, n =* 30 for 30-40s), L1 backgrounds (*n* = 5 for English, *n* = 8 for Chinese, *n* = 8 for Japanese*, n* = 9 for other languages), contexts (*n* = 22 for ESL settings, *n* = 8 for EFL settings), and music experience (*n* = 13 for no, *n* = 17 for yes). According to the results of one-way ANOVAs, the early adulthood group demonstrated significantly lower discrimination scores (greater sensitivity to spectral/temporal details) than the late adulthood group (*p* = .010). The other learner factors did not significantly distinguish the groups’ auditory discrimination and reproduction scores (*p* > .05).

**Table 6.** Summary of One-Way ANOVAs.

| Group variables | Dependent variables | *F* | *p* |
| --- | --- | --- | --- |
| Chronological age (20s vs. 30-40s) | Overall discrimination | 7.720 | .010 |
|  | Overall reproduction | .725 | .402 |
| L1 backgrounds | Overall discrimination | .423 | .738 |
|  | Overall reproduction | 2.073 | .128 |
| Contexts (ESL vs. EFL) | Overall discrimination | .526 | .474 |
|  | Overall reproduction | .045 | .833 |
| Music experience (yes, no) | Overall discrimination | 1.102 | .303 |
|  | Overall reproduction | 2.648 | .115 |

## Discussion

According to our initial test-retest reliability study with 30 L1 and L2 English users, auditory processing abilities could be characterized as two somewhat inter-related but essentially different abilities: (a) encoding spectral and temporal details (formant, pitch, duration, and rise time discrimination), (b) integrating and rhythmic information into motor action (melody and rhythm reproduction). Notably, the results also showed that formant discrimination scores in particular could be considered as a form of composite auditory capacity, as the scores were associated with various aspects of the auditory processing measures, such as temporal acuity (i.e., rise time discrimination) and integration (i.e., rhythm reproduction). The findings here are in line with the view that auditory processing could be a multi-layered phenomenon, characterized by a range of perception and cognitive abilities (Flaugnacco et al., 2014; Surprenant & Watson, 2001; Tierney & Kraus, 2015).

In terms of the test-retest reliability of the discrimination and reproduction tasks, the results of correlation analyses identified medium-to-large correlation coefficients for most of the participants’ discrimination and reproduction test performance between T1 and T2 (*r* = .562-.907). The results suggest that the behavioural measures can reliably tap into various dimensions of participants’ supposedly stable auditory acuity and integration abilities (Moore, 2012). One exception concerns the validity of the duration discrimination test, wherein the participants’ test-rest performance did not reach statistical significance (*p* = .128). This could be ascribed to the possibility that the test format may have failed to capture the particular dimension of the participants’ perception abilities that the test was originally designed to measure (i.e., temporal acuity) or that the sample size was too small (*N* = 30). Given that all the participants took the test online, it is also probable that the duration test needs a more consistent sound system across all participants which can be carefully set up under a laboratory condition. For the last possible scenario, however, it is important to remember that all the other subcomponents of the discrimination tests demonstrated adequate level of test-retest reliability, although they could have been most susceptible to sound system.

Finally, the results of ANOVAs found that participants’ auditory discrimination (but not reproduction) was significantly correlated with their chronological age. The findings concur with the previous literature showing that auditory acuity declines across one’s lifespan (especially after 20s) (Skoe et al., 2015) but that auditory integration is less susceptible to the effects of cognitive aging (Thompson et al., 2015). Interestingly, individual differences in auditory processing were not clearly linked to the other background factors (L1 backgrounds, immersion and music experience). To further examine the biographical correlates of auditory acuity and integration, future studies are called for with larger sample sizes (cf. Saito et al., 2020).

## Future Directions

As we mentioned earlier, the test-retest reliability dataset featured in the current project will continue to expand as our future studies recruit more participants. As such, we aim to provide a more thorough evaluation of the instrument validity in our future projects. To help other L2 acquisition researchers to use the explicit auditory discrimination and reproduction tests, we have two possible scenarios. First, we will develop our HTML webpage so that researchers can easily invite their participants to take the tests, and record/organize their auditory processing scores online. Second, we will provide a PRAAT/Python script, enabling researchers to run the auditory processing test battery as a form of a fixed (but not adaptive) A×B discrimination task (for the same format, see Kempe, Bublitz, & Brooks, 2015).

## Supporting information

Discrimination_Task

Reproduction_Task

